# Selection against archaic hominin genetic variation in regulatory regions

**DOI:** 10.1101/708230

**Authors:** Natalie Telis, Robin Aguilar, Kelley Harris

## Abstract

Traces of archaic hominin DNA persist in the human gene pool, but are systematically depleted around genes and other functionally important genomic regions. This suggests that many Neandertal and Denisovan alleles had harmful effects on hybrid fitness. We hypothesized that if some harmful effects were mediated by gene dysregulation in specific tissues, alleles previously flagged as archaic using a conditional random field (CRF) should be depleted from those tissues’ regulatory enhancers compared to “control” alleles matched for allele frequency and the strength of background selection. By this metric, both Neandertal and Denisovan variation appear depleted from enhancers, particularly enhancers that show pleiotropic activity across tissues. This depletion is driven by young archaic SNPs that the CRF confidently identifies as private to Neandertals or Denisovans; older variants that were likely present in both archaic species are not depleted from enhancers. We found that enhancer pleiotropy is not only a predictor of archaic SNP depletion, but also a predictor of intolerance to new mutations as measured by both phastCons scores and the frequency spectrum of African variation. In other respects, however, the landscape of selection against young archaic alleles appears qualitatively different from the landscape of ordinary purifying selection, suggesting that archaic alleles had a different distribution of fitness effects from ordinary new mutations. Most strikingly, fetal brain and muscle are the tissues most depleted of young archaic variation in their regulatory regions, but only brain enhancers appear commensurately intolerant to new mutations. In contrast, fetal muscle enhancers show no evidence of elevated purifying selection relative to other enhancers. This suggests that epistatic incompatibility between human and archaic alleles is needed to explain the degree of archaic variant depletion from fetal muscle enhancers, perhaps due to divergent selection for higher muscle mass in archaic hominins compared to humans.

## Introduction

Although hybrids between humans and archaic hominins were once viable, fertile and numerous [1, 2, 3, 4], Neandertal and Denisovan alleles have been systematically depleted from the most functionally important regions of the human genome [5, 6, 7]. This pattern implies that archaic introgression often had deleterious consequences for human populations, but it is challenging to fine-map the locations of detrimental archaic alleles and determine the nature of their fitness effects. Petr, et al. recently found that promoters were actually more depleted of introgression than the coding sequences that lie immediately downstream, lending weight to the longstanding hypothesis that gene regulatory mutations underlie much of the functional divergence between closely related lineages of hominins [8, 9, 10]. Two other recent studies found that introgressed alleles are associated with gene expression variation more often than expected by chance [11, 12], implying that even the archaic regulatory variation that remains in the human gene pool is not necessarily benign.

Previous work showed that selection was likely relaxed within the Neandertal exome, leading the accumulation of deleterious mutations, by comparing rates of amino acid change to rates of substitution at synonymous sites [13]. However, it is less straightforward to perform similar analyses on noncoding variantion because of gaps in our understanding of the grammar relating sequence to regulatory function [14, 15]. Allele frequency spectra and patterns of sequence divergence can sometimes provide information about the mode and intensity of selection acting on noncoding regions [16, 17, 18, 19, 20], but introgressed variants have an unusual distribution of allele ages and frequencies that can confound the efficacy of standard methods that assume simple population histories [21]. Reporter assays can directly measure the impact of archaic variants on gene expression *in vitro* [22, 23, 24], but they cannot translate gene expression perturbations into the subtle effects on survival and reproduction that likely determined which archaic variants were purged from human populations.

Petr et al. used a direct *f*_4_ ratio test to conclude that promoters and other conserved noncoding elements harbored less Neandertal DNA than the genome as a whole, but found no similar depletion in enhancers. However, a subsequent study by Silvert et al. came to somewhat different conclusions using different methodology, which involved quantifying the distribution of alleles flagged as likely Neandertal in origin based on their presence in the Altai Neanderthal reference and their absence from an African reference panel [25]. Most such alleles are presently rare (<2% frequency in modern Eurasians), and Silvert, et al. found these rare archaic alleles to be significantly depleted from enhancers. However, archaic variants present at population frequencies of 5% or more were found to occur in enhancers more often than expected by chance. Enhancers containing these common archaic alleles were found to be preferentially active in T cells and mesenchymal cells, perhaps due to positive selection for alleles that alter gene expression in the immune system [25].

The results of these two prior papers are consistent with a model in which most introgressed enhancer sequences have been segregating neutrally within the human gene pool, but in which archaic haplotypes containing private Neandertal enhancer variation were more often selected against than introgressed haplotypes containing private Neandertal variants outside regulatory regions. To interrogate this model more directly, we leverage a set of archaic variant calls that were previously generated using a conditional random field (CRF) approach [5, 7]. The CRF introgression calls are hierarchically organized in a way that correlates with age: some of these alleles are confidently inferred to be either Neandertal or Denisovan in origin, while others might have originated in either archaic species and likely segregated for a longer period of time prior to human secondary contact. We quantify the abundance of young versus old archaic alleles in enhancers as a function of tissue activity, controlling for the amount of background selection enhancers experience, to estimate whether selection likely acted to remove certain classes of archaic variants from regulatory regions.

## Results

### Enhancers appear depleted of Neandertal alleles compared to control regions affected by similar levels of background selection

We intersected the ENCODE Roadmap enhancer calls with the high confidence Neandertal and Denisovan SNP calls generated from the Simons Genome Diversity Panel (SGDP) [7, 26]. Of the two available call sets, we used variant set # 2, which identifies Eurasian variants as Neandertal if they appear more similar to the Altai Neandertal reference than to either an African reference panel or the Altai Denisovan reference. (Similarly, variants are identified as Denisovan if they appear more similar to the Altai Denisovan reference than to an outgroup consisting of Africans plus the Altai Neandertal reference).

Since Neandertal variants in enhancers might have been purged due to linkage disequilibrium with nearby coding variants, we devised a method to measure archaic allele depletion while controlling for the strength of background selection, as quantified by McVicker and Green’s *B* -statistic [27, 5, 7]. We randomly paired each Neandertal variant with two “control” variants matched for both *B* -statistic decile and allele frequency (Figure 1A), then computed the proportion of archaic versus control alleles occurring within enhancers.

**Figure 1:**
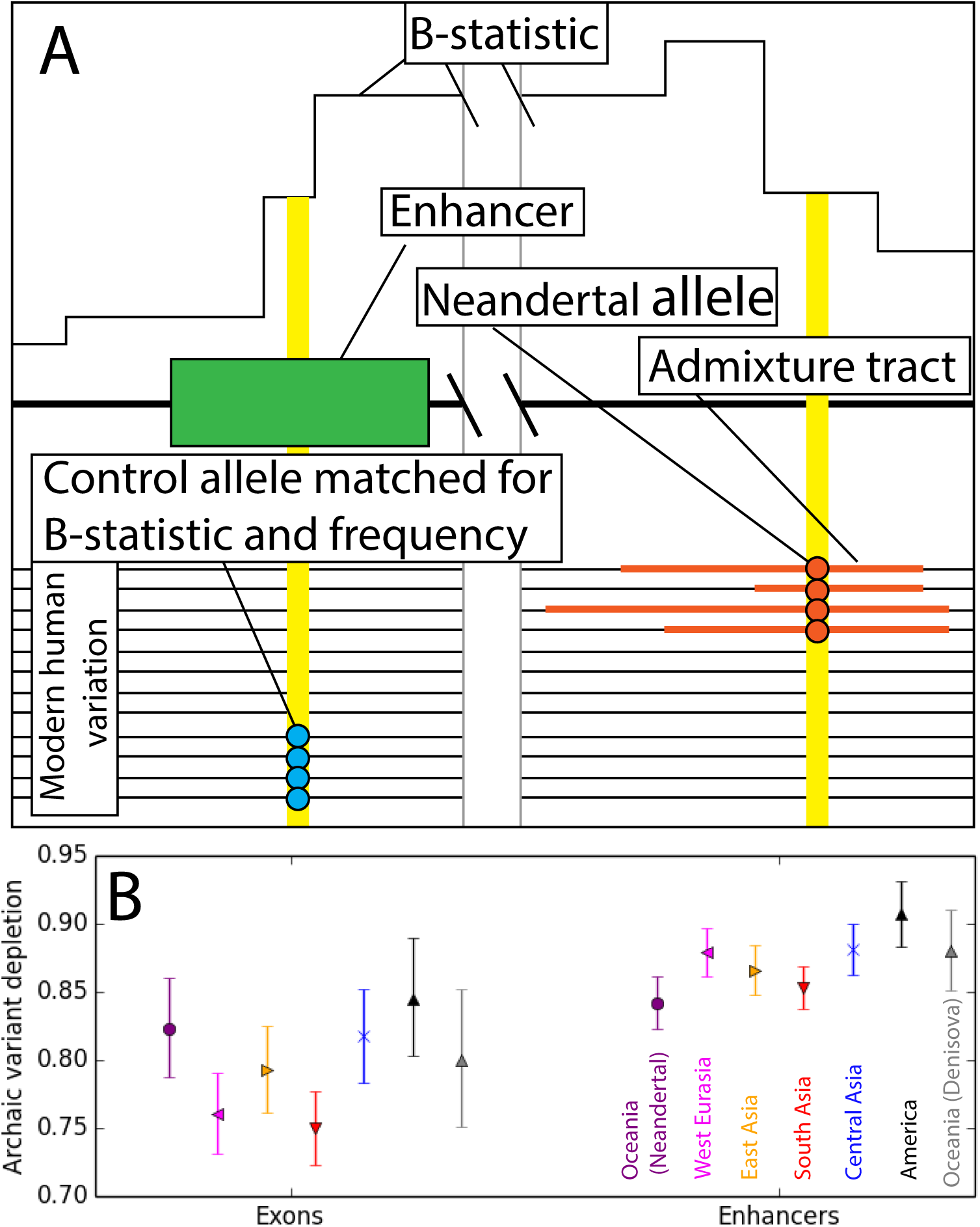
A. This schematic illustrates the process of matching archaic variants to “control” variants with identical allele frequencies and B statistic values. B. Introgressed-to-control variant odds ratios show depletion of Neandertal variants from both exons and enhancers in every population sequenced by the Simons Genomic Diversity Project. In the case of Oceanians, a similar pattern holds for Denisovan variant calls. Error bars span 95% binomial confidence intervals.

In every population, we found that control variants occur within enhancers significantly more often than introgressed variants do (Figure 1B), with depletion odds ratios ranging from 0.84 to 0.91 and 95% binomial confidence intervals excluding an odds ratio of 1. As expected, this method also detects negative selection against introgression in exons. Enard, et al. recently used a related approach to quantify Neandertal introgression in proteins that interact with viruses [28]. To ensure that the linkage structure of the archaic SNPs was not contributing to this result, we sampled an alternate set of controls with a similar linkage block structure; upon substituting these controls for our originally sampled controls, we observed a nearly identical landscape of introgression depletion (Figure S1.1).

### Highly pleiotropic enhancers harbor fewer archaic variants than tissue-specific enhancers

The enhancers annotated by RoadMap exhibit wide variation in tissue specificity. Some are active in only one or two tissues, while others show activity in 20 tissues or more [29]. When we stratified enhancers by pleiotropy number, meaning the number of tissues where the enhancer is active (Figure 2), we found pleiotropy to be correlated with the magnitude of archaic variation depletion (Figure 2).

**Figure 2:**
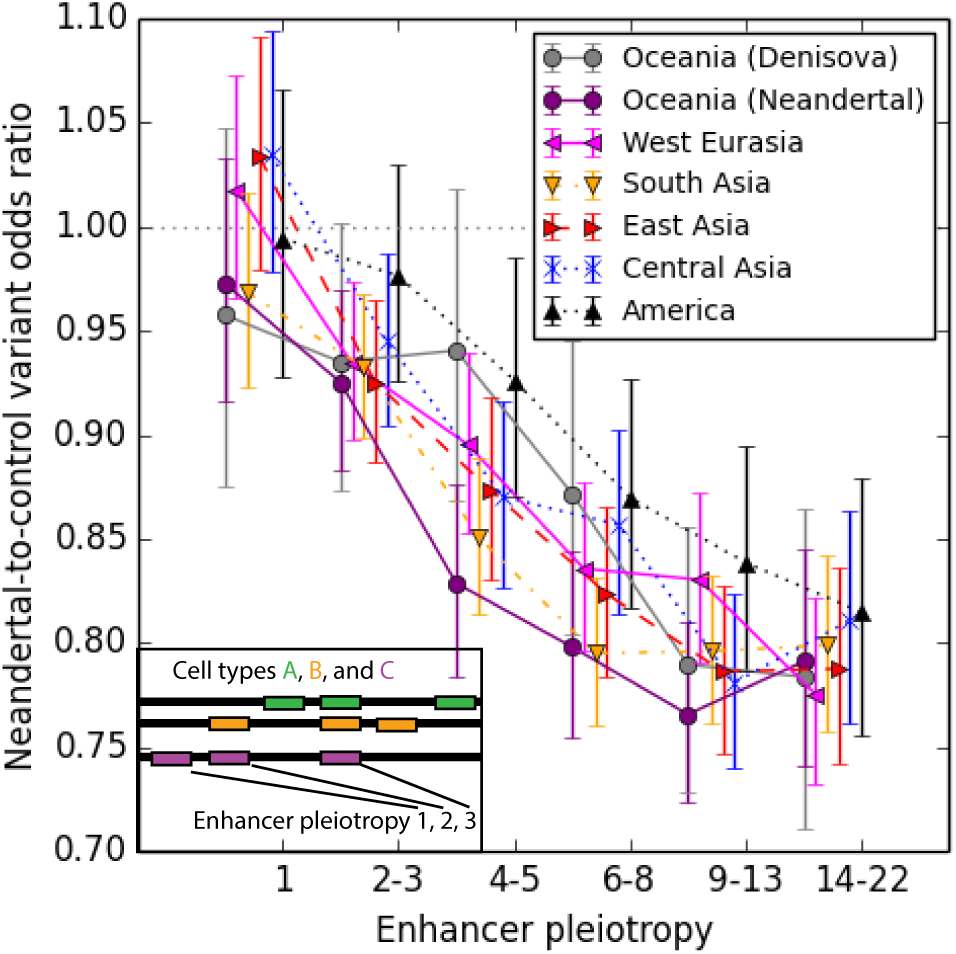
Enhancers active in only a single cell type do not appear depleted of archaic SNPs, whereas enhancers that are active in multiple cell types contain up to 20% fewer archaic variant calls than expected. Error bars span 95% binomial confidence intervals.

If high-pleiotropy enhancers exhibited high sequence similarity between humans, Neandertals, and Denisovans, this could make it hard to detect archaic introgression in these regulatory regions and create the false appearance of selection against introgression. However, we found that the human and archaic reference sequences were actually more divergent in high-pleiotropy enhancers than in other regions (Figure S2.1), making selection against introgression more likely to be responsible for the observed depletion gradient. Enhancer activity is known to increase the mutation rate by inhibiting DNA repair [30, 31], which may explain why highly active enhancers have been diverging between hominid species at an accelerated rate.

We found substantial variation between tissues in the magnitude of archaic SNP depletion (Figure 3A; Figure S3.1), as well as correlation across tissues between depletion of Neandertal variants and depletion of Denisovan variants (*r*^2^ = 0.537, *p* < 4e−5). Enhancers active in fetal muscle, fetal brain, and neurosphere cells are the most strongly depleted of introgressed variation, while enhancers active in fetal blood cells and T-cells, as well as mesenchymal cells, appear the least depleted. We observed no correlation across tissues between the degree of archaic variant depletion and the genetic divergence between archaic and human reference sequences (Figure S3.2). Mesenchymal cells, T-cells, and other blood cells are among the cell types where some adaptively introgressed regulatory variants are thought to be active [32, 33, 34, 25], but our results suggest that selection overall decreased archaic SNP load even within the regulatory networks of these cells. The excess archaic SNP depletion in brain and fetal muscle is a pattern that holds robustly across populations (Supplementary Figure S3.2). Although exons are slightly more depleted of archaic SNPs than enhancers as a whole (see the non-overlapping 95% confidence intervals in Figure 1B), they are actually less depleted of archaic variation than brain or fetal muscle enhancers.

**Figure 3:**
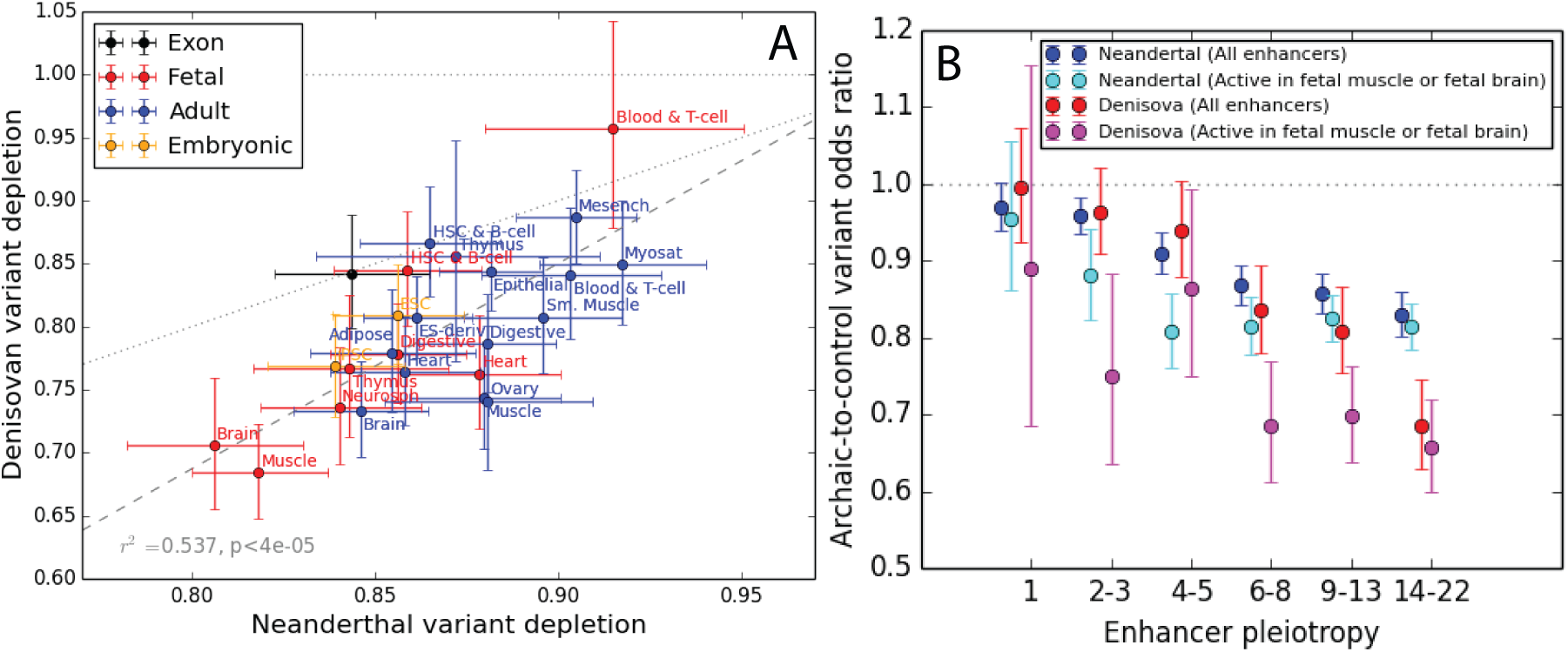
A. Neandertal and Denisovan variant depletion varies between enhancers active in different tissues. Data points that lie below the dotted line correspond to tissues whose enhancers are more depleted of Denisovan SNPs compared to Neandertal SNPs. B. Even after restricting to enhancers active in fetal muscle or fetal brain, the two tissue types most depleted of introgressed variation, pleiotropy remains negatively correlated with archaic SNP depletion. The difference between these two tissues and other tissues is driven mainly by enhancers of intermediate pleiotropy. All error bars span 95% binomial confidence intervals.

Although fetal muscle and fetal brain enhancers are more depleted of archaic SNPs than enhancers active in other tissues, selection acting in these two tissues alone is not sufficient to explain the apparent depletion of archaic SNPs from other tissues’ enhancers. When we computed the magnitude of archaic SNP depletion as a function of pleiotropy in the subset of enhancers that are active in fetal brain, fetal muscle, or both, we still found pleiotropy to be predictive of introgression depletion (Figure 3B).

For enhancers active in 6 or more tissues, depletion of Denisovan variants is notably stronger than depletion of Neandertal variants. One possible culprit is a discrepancy in the power of the CRF to ascertain Neandertal versus Denisovan introgression. The Denisovans who interbred with modern humans were quite genetically differentiated from the Altai Denisovan reference individual, while the Altai Neandertal reference is more modestly divergent from Neandertal introgressed tracts [35], and this likely created differences between species in the sensitivity and specificity of archaic SNP detection.

### Old variation shared by Neandertals and Denisovans was likely less deleterious to humans than variation that arose in these species more recently

To further test the hypothesis that selection acted to purge young, rare archaic variation from enhancers, we leveraged the difference between two introgression call sets that Sankararaman, et al. generated from the SGDP data [7]. As mentioned earlier, we conducted all previous analyses using “Call Set 2,” which was constructed to minimize the misidentification of Neandertal alleles as Denisovan and vice versa. To generate Neandertal Call Set 2, Sankararaman, et al. used an outgroup panel that contained the Altai Denisovan as well as several Yoruban genomes. Similarly, Denisovan Call Set 2 was generated using a panel that included Yorubans plus the Altai Neandertal. In contrast, “Call Set 1” was generated using an outgroup panel composed entirely of Africans, and this procedure identifies more archaic SNPs overall.

Compared to Set 2, we hypothesized that the more inclusive Set 1 calls should contain more old variation that arose in the common ancestral population of Neandertals and Denisovans (Figure 4A). We posited that this older variation might be better tolerated in humans because it rose to high frequency in an ancestral population that was not as divergent from humans as later Neandertal and Denisovan populations. Neandertals and Denisovans also suffered from increasingly severe inbreeding depression as time went on, further increasing the probability that younger variants could have deleterious effects [2, 13].

**Figure 4:**
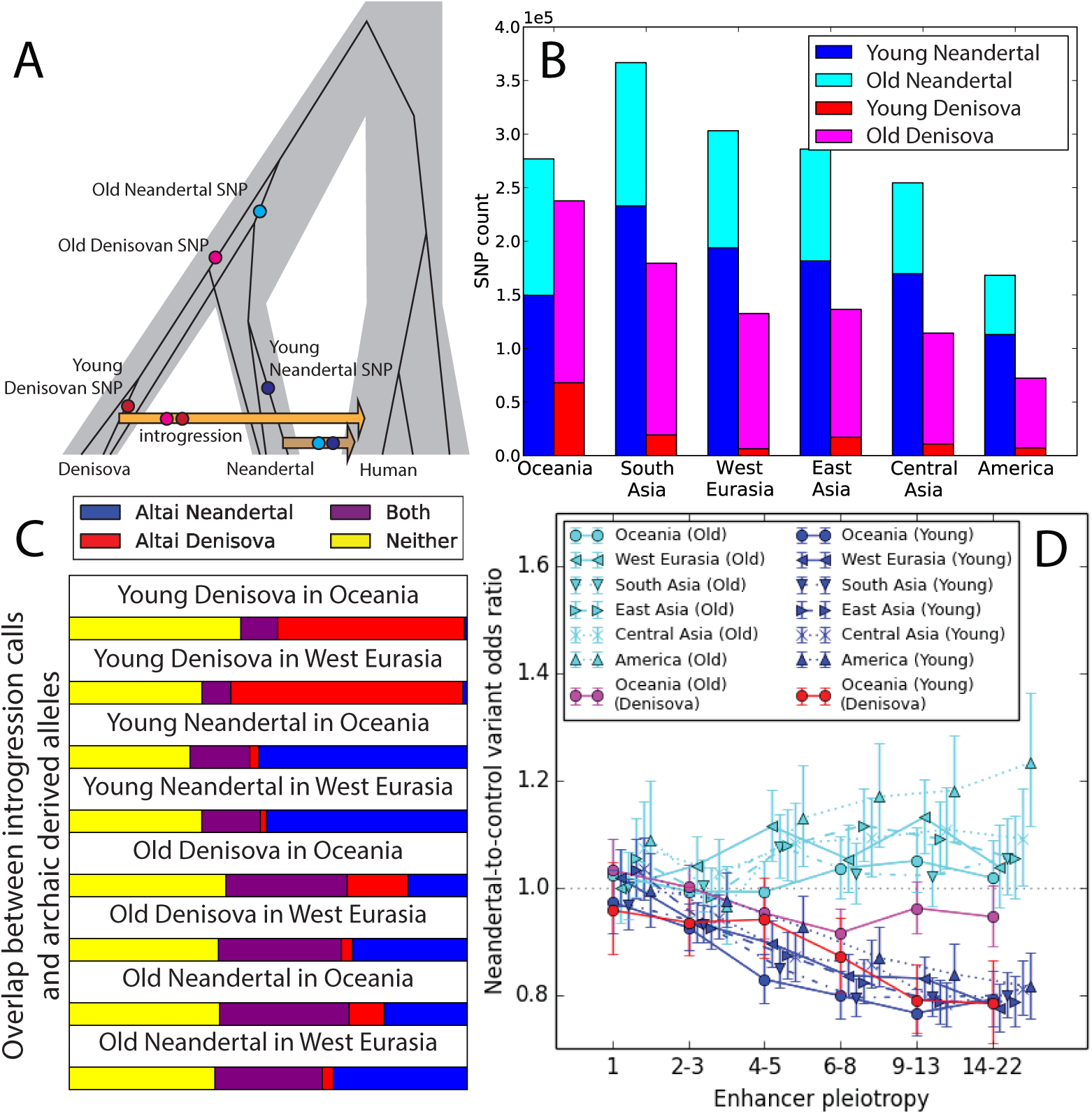
A. We classified introgression calls as “old” or “young” based on their presence in the Sankararaman, et al. Call Sets 1 and 2. By design, the old alleles (present in Call Set 1 but not Call Set 2) are more often shared between Neandertals and Denisovans, and we hypothesize that many of these alleles arose before the Neandertal/Denisovan divergence, as pictured, or else crossed between the two species via Neandertal/Denisovan gene flow. In contrast, we hypothesize that the young alleles most often arose after Neandertals and Denisovans had begun to diverge. B. Numerical counts of old and young introgressed variants in the SGDP human genomes. Young Denisovan variants are likely rare because the Altai Denisovan was not closely related to the Denisovan population that primarily interbred with humans [35]. C. We computed the fraction of CRF introgression calls that occur as derived alleles in the Altai Neandertal genome and/or the Altai Denisovan genome. As expected, old variants are 2-to-4-fold more likely than young variants to occur in both archaic reference genomes. See Supplementary Figures S4.1 and S4.2 for more data on allele sharing between introgression calls and the reference archaic genomes. D. In contrast to the young archaic variation considered elsewhere in this paper, old archaic variation is not measurably depleted from enhancers, even enhancers active in numerous tissues. All error bars span 95% binomial confidence intervals.

To test our hypothesis that Set 2 might be enriched for deleterious variation, we compiled sets of “old” Neandertal and Denisovan variation comprised of their respective Set 1 introgression calls and excluding all Set 2 introgression calls. In every population, young Neandertal variants outnumber old Neandertal variants, but conversely, old Denisovan variants outnumber young Denisovan variants (Figures 4B, S4.2). The CRF may have been better powered to detect young Neandertal variants compared to young Denisovan variants due to the aforementioned closer relationship of the reference Neandertal to archaic individuals who interbred with humans [35]. As expected, old variants are more likely than young variants to be present in both archaic reference genomes rather than just one, though more than 30% of calls in each category are absent from both archaic references and are presumably identified as archaic due to patterns of linkage disequilibrium (Figure 4C).

In contrast to the young Set 2 introgression calls, old calls are not measurably depleted from enhancers compared to control variants matched for allele frequency and B statistic (Figure 4D). This suggests that the introgression landscape was shaped mainly by selection against Neandertal and Denisovan variants that arose relatively close to the time that gene flow occurred, not variation that arose soon after their isolation from humans. Many populations actually show a slight enrichment of old archaic variants in enhancers compared to controls, as shown in Figure 4D by 95% confidence intervals that exclude an odds ratio of 1. These sets of old, shared variants are possibly enriched for beneficial alleles that swept to high frequency in the common ancestor of Neandertals and Denisovans. They should at least be depleted of deleterious variation compared to our control alleles that likely arose more recently in humans. In several cases, the odds ratio enrichment of old archaic variation in enhancers actually trends upward with increasing pleiotropy, possibly because the highest-pleiotropy enhancers show the most divergence between human and archaic reference genomes (Figure S2.1). If archaic SNPs in enhancers were generally neutral or selectively favored, the positive correlation between pleiotropy and human/archaic divergence would lead us to predict the odds ratio trend that is observed for old archaic variants, not the opposite correlation with pleiotropy that is observed for young archaic variants.

Neandertals and Denisovans are thought to have begun diverging about 640,000 years ago [36]. Since this is long enough to efficiently purge deleterious variation, any surviving archaic variation that predates this split is likely to have nearly neutral or beneficial fitness effects, assuming no negative epistasis with human variation. We can see this from a simple population genetic calculation: assuming that the Neandertal/Denisovan effective population size was about 4,000 [2] and their generation time is 30 years, 4*Ne* for these species would be 480,000. This implies that more than half of the variation that segregated neutrally in the ancestral Neandertal/Denisovan population would have been fixed or lost by the time the two species interbred with humans, leaving ample time for deleterious ancestral variation to be purged. 480,000 years also predates the estimate of the start of the bottlenecks that affected Neandertals and Denisovans [2], so variation that predates this period may have been efficiently purged of deleterious alleles that would have segregated neutrally if they had arisen after the start of the bottleneck period. Some old variants might be younger than the Neandertal/Denisova split if they crossed between the boundaries of these species by introgression; Neandertals and Denisovans are known to have interbred with each other while still maintaining distinct gene pools. This population history suggests that gene flow between Neandertals and Denisovans may be enriched for variants that are benign on a variety of genetic backgrounds [37], making them more likely to be benign on a human background as well.

### Introgressed variants and recent mutations have been differently selected against as a function of enhancer activity

We next investigated whether the enhancers most depleted of young archaic variants are simply the enhancers most intolerant to new mutations, leveraging the fact that natural selection allows neutral and beneficial mutations to reach high frequencies more often than deleterious mutations do [38, 39] (Figure 5A). Working with the site frequency spectrum (SFS) of African enhancer variation from the 1000 Genomes Project, we computed the proportion of variants segregating in enhancers that are singletons and compared this to the proportion of singletons in the immediately upstream enhancer-sized regions (Figure 5B). Neandertals contributed much less genetic material to sub-Saharan Africans compared to non-Africans [1, 4], meaning that Neandertal alleles should have little direct effect on the African site frequency spectrum.

**Figure 5:**
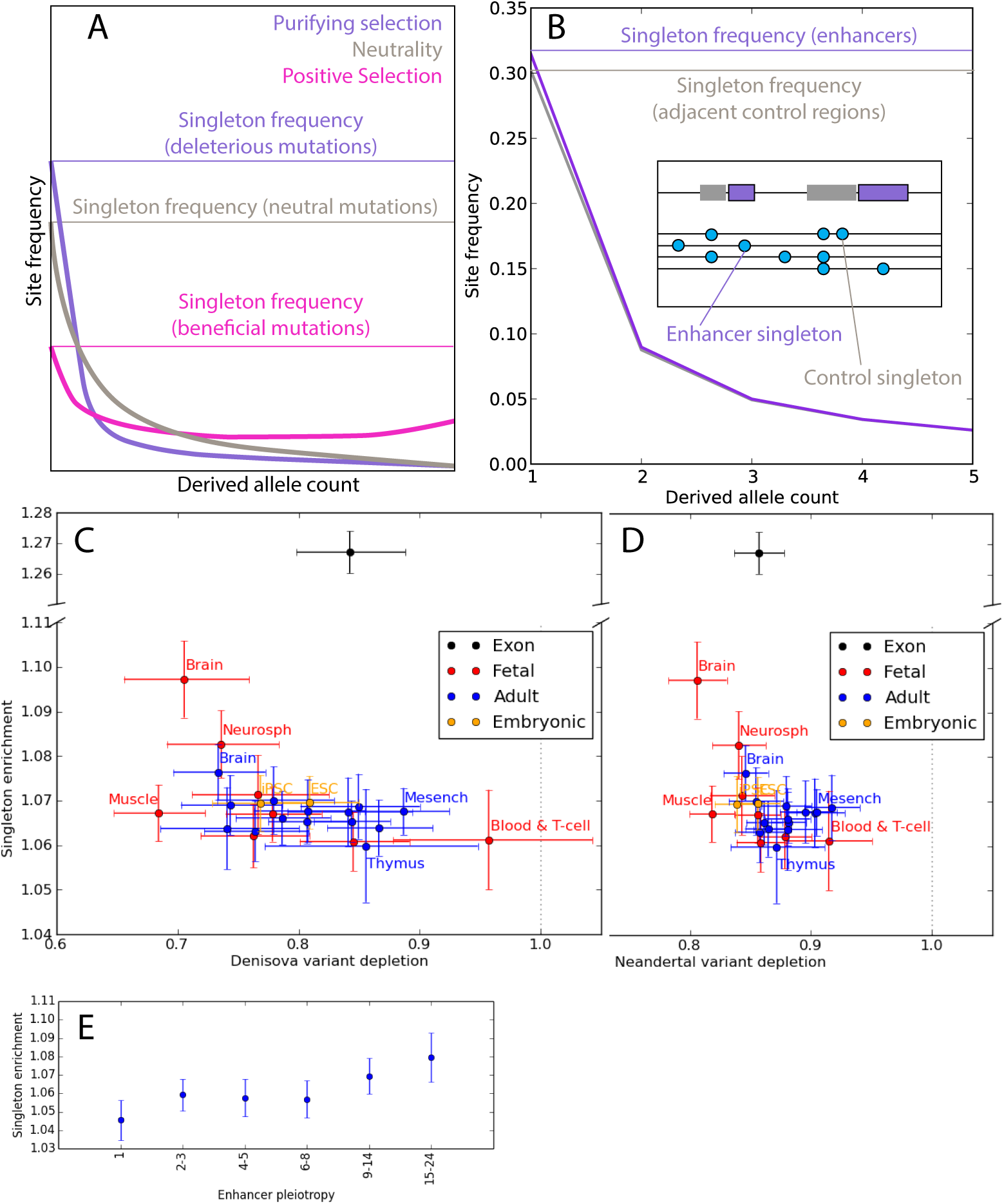
A. Theory predicts that the site frequency spectrum (SFS) becomes skewed toward rare variants by the action of purifying selection. B. In African data from the 1000 Genomes project, the enhancer SFS has a higher proportion of singletons compared to control regions adjacent to enhancers. C. Every tissue type’s enhancer complement is enriched for singletons compared to adjacent control regions. This comparison of singleton enrichment odds ratios to Denisovan depletion odds ratios shows that fetal brain, neurosphere cells, and adult brain are outliers under stronger constraint. The y axis has been split to accommodate the magnitude of singleton enrichment in exons. Error bars span 2 binomial test standard errors. D. Comparison of the singleton enrichment landscape to the Neandertal depletion landscape. E. Enhancer pleiotrophy is negatively correlated with singleton enrichment, though even enhancers of pleiotropy 1 have a singleton enrichment odds ratio significantly greater than 1. All error bars span 95% binomial confidence intervals.

One caveat is that this strategy will not detect the effects of strongly deleterious mutations that do not segregate long enough to affect the frequency spectrum’s shape. However, strongly deleterious mutations are not expected to contribute to mutation load differences between populations, making it appropriate to focus on identifying regions whose variation is affected by selection against weakly deleterious mutations.

By comparing enhancers to immediately adjacent regions, we control for the potentially confounding effects of recombination rate, background selection, and sequencing read depth. Although enhancers likely have elevated mutation rates because transcription factor binding impairs DNA repair [30], the proportion of variants that are singletons is independent of mutation rate as long as the mutation rate has remained constant over time [40]. Enhancers admittedly have higher GC content than adjacent control regions, but the enrichment of singletons in the enhancer SFS holds separately for SNPs with AT ancestral alleles and SNPs with GC ancestral alleles (Figure S5.1). This suggests that the SFS is not enriched for singletons because of a force like biased gene conversion, which only depresses the frequencies of mutations from GC to AT and instead increases the frequencies of mutations from AT to GC [41].There is also no apparent correlation across tissues between GC content and the enrichment of rare variants in enhancers (Supplementary Figure S5.2). We conclude that purifying selection is likely driving the difference between the SFSs of enhancers and control regions, not base composition or biased gene conversion.

Although enhancers broadly show evidence of purifying selection against both archaic variation and new mutations, the strength of selection against these two types of perturbation is poorly correlated among tissues (Figure 5C, D). Although singleton enrichment appears nominally correlated with Neandertal depletion (*r*^2^ = 0.31, *p* < 0.004) and Denisovan depletion (*r*^2^ = 0.27, *p* < 0.009), this correlation disappears when brain tissues are excluded (Neandertal *p* < 0.42; Denisovan *p* < 0.10; see Figure S5.3). Fetal brain, neurosphere cells, and, to a lesser extent, adult brain are the tissues whose active enhancers show the most singleton enrichment, suggesting that mutations perturbing brain development have an outsize probability of deleterious consequences. In contrast, fetal muscle enhancers show no evidence of unusual selective constraint despite their strong depletion of both Neandertal and Denisovan ancestry. We obtain categorically similar results when we estimate selective constraint using phastCons scores rather than singleton enrichment (Figure S5.4).

Enhancer pleiotropy is positively correlated with singleton enrichment as well as the depletion of archaic alleles (Figure 5E). This observation may be related to experimental evidence that the most highly pleiotropic enhancers tend to have the most consistently conserved functioning across species [42]. One difference, however, is that enhancers active in only a single tissue (pleiotropy number 1) still show significant evidence of selection against new mutations despite their lack of any evidence for selection against archaic introgression (O.R. 95% C.I. excludes 1).

## Discussion

Most methods for identifying introgressed archaic haplotypes rely on putatively unadmixed outgroup data. Chen, et al. recently showed that use of an African outgroup can confound measurements of introgression fraction differences between populations, causing less introgression to be detected in Europeans compared to Asians because Europeans exchanged more recent migrants with Africa [4]. Our analysis of young versus old CRF-based calls shows that the choice of outgroup can also affect the distribution of archaic allele calls across functional versus putatively neutrally evolving genomic regions. This implies that outgroup panel use can interfere with efforts to estimate unbiased Neandertal and Denisovan admixture fractions, but does not imply that unbiased admixture fractions are necessarily the most powerful statistic for detecting the footprints of selection against archaic alleles. The subset of archaic haplotypes that are most divergent from outgroup panels are by definition enriched for mutations that may have detectable fitness effects, whereas archaic haplotypes that are less divergent and more difficult to detect computationally are more likely to segregate neutrally in human populations. In reaching such conclusions, proper care must be taken to control for rates of human/archaic reference divergence, which can vary across the genome. In enhancers, however, we found archaic/human divergence to be elevated, which likely enhanced the power of the CRF to discover introgression overlapping these regions. This suggests that selection is needed to explain the observed depletion of young archaic variants from enhancers.

Two sources of dysfunction are thought to drive selection against archaic introgression: excess deleterious mutation load in inbred Neandertal and Denisovan populations [43, 44] and accumulation of epistatic incompatibilities due to divergent selective landscapes [5, 7, 45]. Both forces have the potential to affect enhancers, and our results confer some ability to distinguish between the two. In particular, the weakness of the correlation between archaic allele depletion and singleton enrichment furnishes useful insights into the fitness effect differences between *de novo* human mutations and young introgressed archaic alleles. This difference appears starkest when comparing enhancers to exons, which are known to evolve more slowly than enhancers over phylogenetic timescales [46, 47, 48], implying that selection acts more strongly against new coding mutations compared to new regulatory mutations. However, despite their different levels of selective constraint against new mutations, exons and enhancers show evidence for selection against archaic alleles (Figure 5C, D), suggesting that regulatory effects may have played a significant role in shaping the landscape of Neandertal and Denisovan introgression.

When a set of regulatory elements is more depleted of introgression than expected given their level of selective constraint, this suggests that the Neandertal and Denisovan selective landscape may have diverged from the human one in these regions. Fetal muscle enhancers appear to fit this profile, with unremarkable singleton enrichment and phastCons scores but strong depletion of young archaic variants. Archaeological evidence indicates that Neandertals had higher muscle mass, strength, and anatomical robustness compared to humans [49, 50], supporting the idea that the two species had different fetal muscle growth optima. We have no direct knowledge of Denisovan muscle anatomy, but the depletion of Denisovan DNA from muscle enhancers may suggest that they shared Neandertals’ robust phenotype, assuming that phenotype is mediated by gene regulation in fetal muscle.

In contrast to muscle, mutation load is a more attractive candidate cause for the depletion of archaic alleles from brain enhancers. Our conclusion that brain enhancers experience high deleterious mutation rates is bolstered by prior knowledge of many *de novo* mutations in these regions that cause severe developmental disorders [51, 52, 53].

Both genetic load and hybrid incompatibilities might drive the correlation we have found between enhancer pleiotropy and archaic allele depletion. Steinrücken, et al. noted that epistatic incompatibilities are most likely to arise in genes with many interaction partners; when a gene is active in multiple tissues, it must function as part of a different expression network in each tissue, which could create additional constraints on enhancers that must coordinate expression correctly in several different contexts. Our results thus imply that introgression is most depleted from enhancers that must function within a variety of cell-specific regulatory networks. We also know that genes expressed in many tissues evolve more slowly than genes expressed in few tissues because they have greater potential for functional tradeoffs [55, 10], and a mutation that disrupts the balance of a functional tradeoff is likely to have a deleterious effect. This idea is corroborated by our finding that pleiotropic enhancers are more constrained. One caveat is that highly pleiotropy enhancers may be the easiest to experimentally identify. If the RoadMap call sets of tissue-specific enhancers contain a higher proportion of false positives, this might inflate our estimate of the correlation between pleiotropy and selective constraint.

Both genetic load and epistatic incompatibilities are expected to “snowball” over time, making young archaic variation more likely to be deleterious in hybrids compared to older, high frequency archaic variation. Part of this effect might be due to positive selection on beneficial introgressed alleles that have risen to high frequency in multiple populations. As more methods for inferring admixture tracts are developed, our results underscore the importance of investigating how they might be biased toward young or old archaic variation and using this information to update our understanding of how selection shapes introgression landscapes. Regulatory mutations appear to have created incompatibilities between many species that are already in the advanced stages of reproductive isolation [56, 57, 58, 59], and our results suggest that they also harmed the fitness of human/Neandertal hybrids during the relatively early stages of speciation between these hominids. As more introgression maps and functional genomic data are generated for hybridizing populations of non-model organisms, it should be possible to measure the prevalence of weak regulatory incompatibility in more systems that exist in the early stages of reproductive isolation and test how many of the patterns observed in this study occur repeatedly outside the hominoid speciation continuum.

## Acknowledgements

We are grateful to Jonathan Pritchard, Sriram Sankararaman, Josh Schraiber, and members of the Harris Lab for helpful discussions, and we thank Rasmus Nielsen for manuscript comments. We also thank three anonymous reviewers whose comments greatly improved the manuscript. We acknowledge financial support from the following grants awarded to K.H: NIH grant 1R35GM133428-01, a Burroughs Wellcome Fund Career Award at the Scientific Interface, a Searle Scholarship, a Sloan Research Fellowship, and a Pew Biomedical Scholarship.

## Data Availability Statement

All datasets analyzed here are publicly available at the following websites:

|  |  |
| --- | --- |
| CRF Introgression Calls | https://sriramlab.cass.idre.ucla.edu/<br>public/sankararaman.curbio.2016/summaries.tgz |
| SGDP | https://www.simonsfoundation.org/<br>simons-genome-diversity-project/ |
| RoadMap | https://personal.broadinstitute.org/meuleman/reg2map/<br>HoneyBadger2_release/ |
| 1000 Genomes Phase 3 | http://www.1000genomes.org/category/phase-3/ |

## Code availability statement

Summary data files and custom python scripts for reproducing the paper’s main figures are available at https://github.com/kelleyharris/hominin-enhancers/.

## Author contribution statement

N.T. and K.H. conceived and designed the project. N.T., R.A., and K.H. performed the analyses. K.H. wrote the paper.

## Methods

### Extraction of Neandertal and Denisovan variant sets

Neandertal and Denisovan variant call sets were downloaded from https://sriramlab.cass.idre.ucla.edu/public/sankararaman.curbio.2016/summaries.tgz. These files classify a haplotype as archaic if it is classified as archaic with ≥ 50% probability by the conditional random field analyses reported in [7]. Using these summaries, we classify a variant as archaic if 100% of the haplotypes on which it appears are classified as such. Unless otherwise stated, all Neandertal and Denisovan variants are obtained from the respective summary call set “2,” which we refer to in the text as the proximal call sets. To construct the distal Neandertal call set analyzed in Figure 4, we included all variants from Neandertal Set 1 except any variants that also appeared in Neandertal Set 1 or Denisovan Set 1. Similarly, the distal Denisovan call set included all variants present in Denisovan Set 1 except those variants also present in Neandertal Set 2 or Denisovan Set 2. Chromosome X was excluded given its unique systematic depletion of Neandertal and Denisovan variants. Across SGDP populations, the number of SNPs identified as Neandertal in origin ranges from 109,253 (in West Eurasia) to 233,013 (in South Asia). The number of introgressed Denisova SNPs ranges from 6,437 (in West Eurasia) to 68,061 (in Oceania).

### Classifying enhancers by tissue type and pleiotropy number

Cell lines were classified into tissue types using the tissue assignment labels from the July 2013 RoadMap data compendium, available at https://personal.broadinstitute.org/meuleman/reg2map/HoneyBadger2_release/DNase/p10/enh/state_calls.RData. Whenever a tissue type contained both fetal and adult cell lines, we further subdivided that tissue type into “Adult” and “Fetal.” We then computed a pleiotropy number for each enhancer by counting the number of distinct tissue type labels in the cell lines where that enhancer is annotated as active. Three separate states are used to denote enhancer activity in the honey badger model: states 6, 7, and 12 denote genic enhancers, enhancers, and bivalent enhancers, respectively, and we could each of these states as equivalent evidence of enhancer activity. Fetal and adult tissue types are counted as distinct tissues for the purpose of this computation.

### Testing for depletion of archaic variation relative to matched control variation

To estimate the strength of background selection experienced by human genomic loci, B-statistic values ranging on a scale from 1 to 1000 were downloaded at http://www.phrap.org/software_dir/mcvicker_dir/bkgd.tar.gz. We quantized these values by rounding them down to the nearest multiple of 50 B-statistic units, then lifted them over from hg18 coordinates to hg19 coordinates. Each SNP in the SGDP data was assigned the B statistic value of the closest site annotated by McVicker, et al.

Our tests for depletion of archaic variation are computed relative to non-archaic control SNPs that have the same joint distribution of allele frequency and B statistic as the SNPs annotated as archaic in origin. See the next section, “Detailed sampling procedure for matched control SNPs”, for more information on how these matched control sets are obtained.

Assume that 𝒜 is a set of *A* archaic SNPs and 𝒞 is a set of 2 × *A* matched controls. (We chose to sample 2*A* controls rather than *A* controls to reduce the stochasticity of the control set and decrease the size of the confidence intervals on all computed odds ratios). To test whether archaic variation of this type is enriched or depleted in a set 𝒢 of genomic regions, we start by counting the number *A*_*G*_ of archaic SNPs contained in 𝒢 and the number *C*_*G*_ of control SNPs contained in 𝒢. We say that this type of archaic variation is depleted from 𝒢 if the odds ratio (*A*_*G*_*/*(*A* − *A*_*G*_))*/*(*C*_*g*_*/*(2*A* − *C*_*G*_)) is less than 1.

To assess the significance of any enrichment or depletion we measure, we ask whether the corresponding log odds ratio log(*A*_*G*_) + log(2*A* − *C*_*G*_) − log(*A* − *A*_*G*_) − log(*C*_*G*_) is more than two standard errors away from zero. The standard error of this log odds ratio is 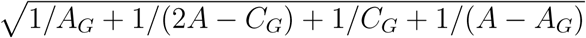. In each forest plot presented in the manuscript, this formula was used to draw error bars that span two standard errors in each direction.

### Detailed sampling procedure for matched control SNPs

For each archaic SNP set (Neandertal 1, Neandertal 2, Denisovan 1, and Denisovan 2) and each population *p*, we counted the number *A*_*p*_(*b, c*) of alleles with B-statistic value *b* and derived allele count *c* in population *p*, counting the allele as archaic if all derived alleles were annotated as present on archaic haplotypes in the relevant call set of population *p*. We then counted the number *N*_*p*_(*b, c*) of non-archaic alleles with B-statistic *b* and derived allele count *c*. In order for a SNP to count as non-archaic, none of its derived alleles could be present on a haplotype from population *p* that was called as archaic in either call set 1 or call set 2. A set 𝒞_*p*_(*b, c*) of 2 × *A*_*p*_(*b, c*) control SNPs was then sampled uniformly at random without replacement from the *N*_*p*_(*b, c*) control candidate SNPs. In the rare event that *N*_*p*_(*b, c*) < 2 × *A*_*p*_(*b, c*), the control set was defined to be the entire set *N*_*p*_(*b, c*) and an extra 2 × *A*_*p*_(*b, c*) − *N*_*p*_(*b, c*) SNPs from the control set were chosen unformly at random to be counted twice in all analyses.

Several analyses in the paper are performed on a merged set of archaic variation compiled across populations. To form the archaic SNP set 𝒜(*b, c*), we merged together the archaic SNP sets 𝒜_*p*_(*b, c*) across all populations *p*. For each site where the derived allele was present in two or more populations, it was randomly assigned one population of origin. This population assignment process yielded new archaic allele counts 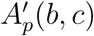 that might be less than the counts *A*_*p*_(*b, c*) due to the deletion of duplicate SNPs. For each population *p*, we sampled 2 × 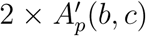 control SNPs from population *p* as before and merged all of these control sets together to obtain a merged control set 𝒞(*b, c*). In the unlikely event that a single control allele is sampled in two or more populations, this control SNP will simply be counted two or more times during downstream analyses.

To obtain sets of distal archaic SNPs and controls, we must be careful about how we subtract call set 2 from call set 1. We want to sample control SNPs such that no control SNP is part of call set 2 for any archaic species in any population. To achieve this, the set of distal Denisovan SNPs 𝒜^(*D*1−2)^(*b, c*) is defined as the set of all SNPs that are present in Denisovan call set 1 𝒜^(*D*1)^(*b, c*) but absent from both the Denisovan call set 2 𝒜^(*D*2)^(*b, c*) and the Neandertal call set 2 𝒜^(*N* 2)^(*b, c*). To generate the corresponding control set 𝒞^(*D*1−2)^(*b, c*), we first look within each population to generate the superset of matched control SNPs 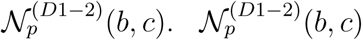 is defined as the set of all SNPs present in population *p* in Denisovan call set 1 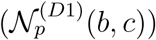 but absent from the population-merged sets of nonarchaic variants from Neandertal Set 2 (𝒩 ^(*N* 2)^(*b, c*)) plus Denisovan Set 2 (𝒩 ^(*D*2)^(*b, c*)). Once we have the population-specific candidate control sets 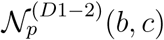, we randomly assign each archaic SNP from 𝒜^(*D*1−2)^(*b, c*) to one of the populations where the derived allele is called as archaic, obtaining population-specific call sets 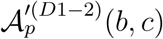 that each contain 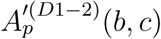 SNPs. As described earlier, we sample 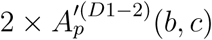 control SNPs uniformly at random from each set 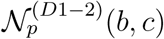 and merge these control sets together to obtain a merged set of distal controls 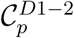.

### Sampling an alternate set of controls to approximate the clustering and LD structure of introgressed variation

The archaic variants present in the human population do not have independent demographic and selective histories, but are in many cases organized into linked archaic haplotypes. To measure whether this LD structure might affect the apparent depletion of archaic alleles from enhancers, we sampled an alternate set of control SNPs whose LD structure is more similar to the LD structure of the introgressed SNPs. LD has the effect of organizing introgressed SNPs into clusters of close-together variants that have similar allele frequencies, and such clustering could increase the probability that a short enhancer sequence might fall into a gap between introgressed SNPs.

To enable sampling of control SNPs in a way that matches the clustering of archaic SNPs, we first organized the introgressed SNPs into blocks, considering two SNPs to be part of the same block if they are less than 20 kb apart and have minor allele counts that differ by at most 1. After organizing the archaic SNPs into these blocks, we counted blocks of control SNPs from the same population VCF that had approximately the same allele frequency and B statistic value. In order to find enough matched control blocks, we relaxed the assumption that archaic and control SNPs should have exactly the same allele frequency and B statistic. Instead, we binned minor allele count into log-2 spaced bins (minor allele count 1, 2, 3-4, 5-8, 9-16, 17+) and required each control SNPs cluster to match the minor allele count bins of the matched archaic cluster. Specifically, given a block of *k* archaic SNPs that we infer to be a haplotype block, we assign the minor allele count bin of that block to be the most common bin occupied by the *k* SNPs. We assigned the B statistic of the block to be the median B statistic of the *k* SNPs. We then counted the number of blocks of *k* consecutive non-introgressed SNPs that had the same minor allele count (plus or minus 1), the same median B statistic, and whose genomic span in base pairs was within a factor of 2 of the span in base pairs of the archaic SNP set. We selected one of these blocks uniformly at random to be the control SNP block matched to the archaic SNP block.

### Quantifying singleton enrichment in the 1000 Genomes site frequency spectrum

Let 𝒢 be a set of enhancers or other genomic regions. To test whether 𝒢 is under stronger purifying selection than its immediate genomic neighborhood, we compared its site frequency spectrum (SFS) to the SFS of a region set 𝒢′defined as follows: 𝒢 can always be defined as a collection of genomic intervals 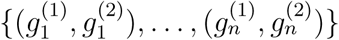, where each 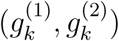 is a pair of genomic coordinates delineating a piece of DNA contained entirely within the set 𝒢. We define 𝒢′ to be the collection of genomic intervals 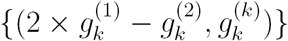, i.e. the set of intervals immediately adjacent on the left to the intervals that make up 𝒢. (We are slightly abusing notation here by failing to note that different chromosomes have different coordinate systems).

We computed folded site frequency spectra for 𝒢 and 𝒢′using the African individuals in the 1000 Genomes Phase 3 VCF, excluding SNPs that do not pass the VCF’s default quality filter. Let *S*_*G*_ and *S*_*G*′_ be the numbers of singletons that fall in into the regions 𝒢 and 𝒢 ′, respectively, and let *N*_*G*_ and *N*_*G*′_ be the numbers of non-singleton variants that fall into these regions. We say that *G* is enriched for singletons if the odds ratio (*S*_*G*_*/N*_*G*_)*/*(*S*_*G*′_ */N*_*G*′_) is greater than 1. To assess the significance of any enrichment or depletion, we use the fact that the standard error of this binomial test is 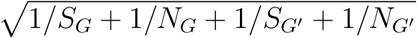. All singleton enrichment plots in this manuscript contain error bars that span 2 standard errors above and below the estimated odds ratio.

**Figure S1.1.**
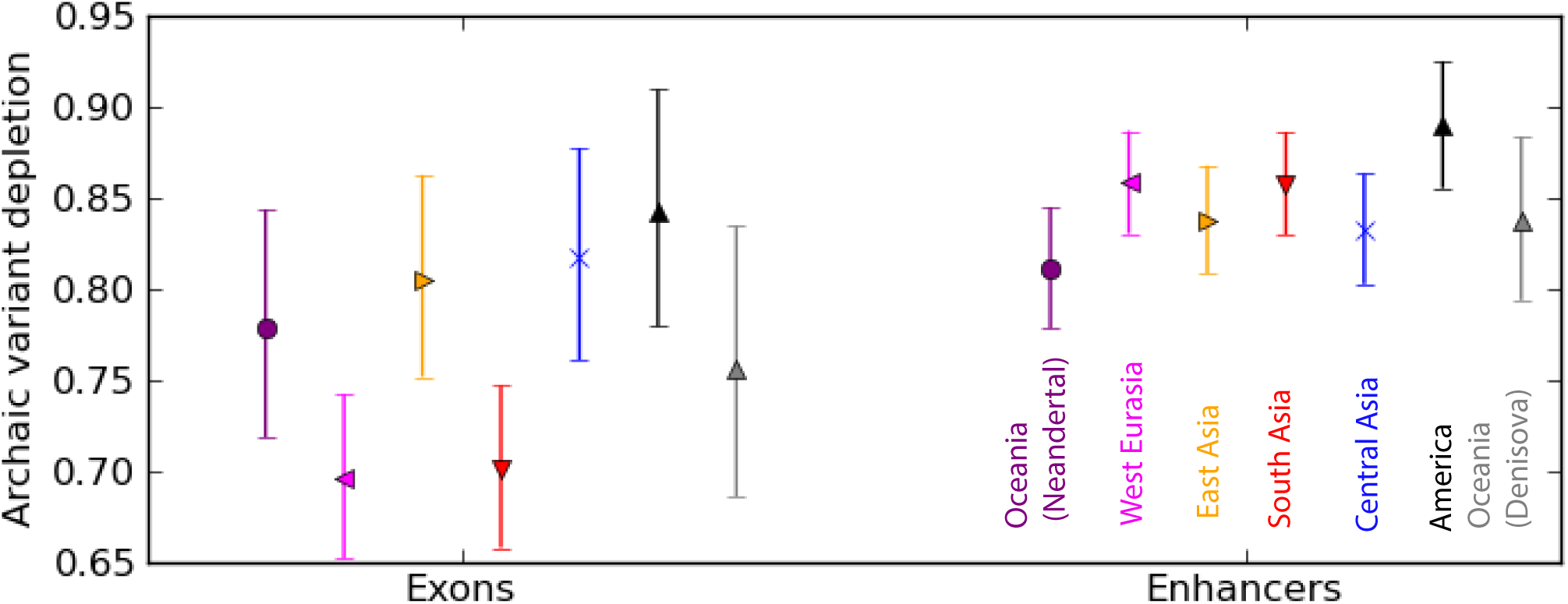
We replicated the analysis from Figure 1B using archaic SNPs and controls sampled to match the clustering induced by LD structure. The depletion of archaic SNPs from exon and enhancers is nearly identical to the depletion measured using controls not sampled to match the spatial clustering of introgressed SNPs.

**Figure S2.1.**
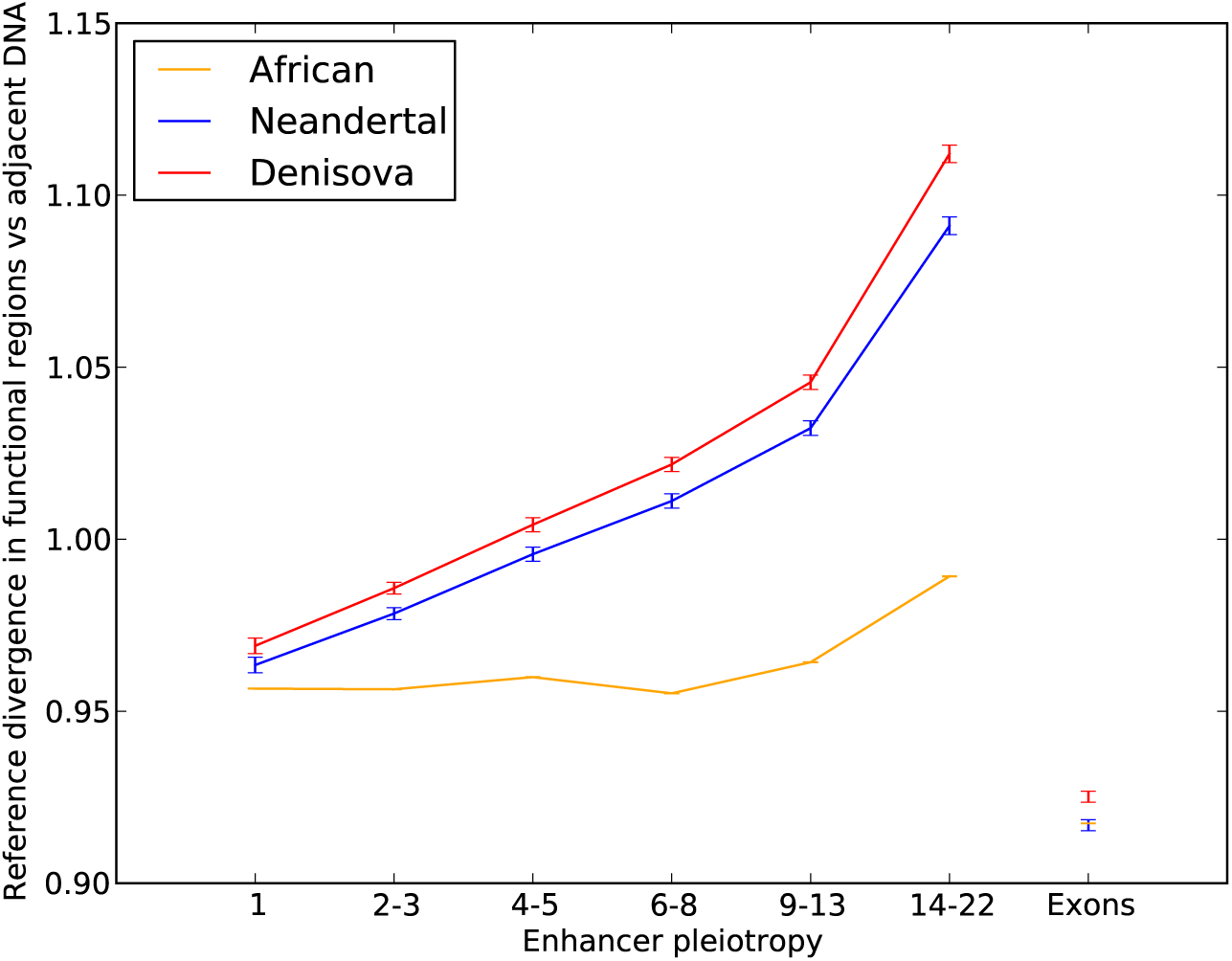
If human and archaic genomes were less diverged from each other in high-pleiotropy enhancers than within other regions of the genome, this could in theory explain why introgressed archaic SNPs are depleted from high pleiotropy enhancers. However this plot shows that the opposite pattern is true: human divergence from reference archaic genomes is higher within high-pleiotropy enhancers than within matched sets of enhancer-sized control regions located 3’-adjacent to enhancers in the genome. To see this, we let *π*_Nean_(*p*) be the pairwise divergence between a Neandertal haplotype (averaged between the two haplotypes of the Altai Neandertal reference) and the human reference genome measured across the set of enhancers of pleiotropy *p*, and let 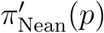 be the pairwise divergence between Neandertal and human within the adjacent matched control regions. The line labeled “Neandertal” in the above plot shows 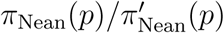 as a function of pleiotropy number *p*. We also measured the average divergence of the human reference genome from an Altai Denisovan reference haplotype and from the set of African genomes sequenced as part of the 1000 Genomes project. The results, 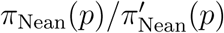, 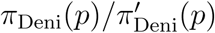, and 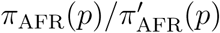, are plotted together with error bars derived from the binomial approximation to the Bernoulli distribution. Divergence between each pair of genomes is consistently lower in exons compared to adjacent control regions, an observation that is consistent with the conserved nature of exonic sequence. In contrast, enhancers are overall more diverged between populations and species than control regions are, and this acceleration of divergence is positively correlated with enhancer pleiotropy. This correlation is stronger for archaic vs human comparisons than for the African vs human reference genome comparison. In the absence of selection against Neandertal DNA in enhancers, this enrichment of divergence should cause a pattern of archaic SNP enrichment within high-pleiotropy enhancers, not the pattern of depletion that we in fact observe.

**Figure S2.2.**
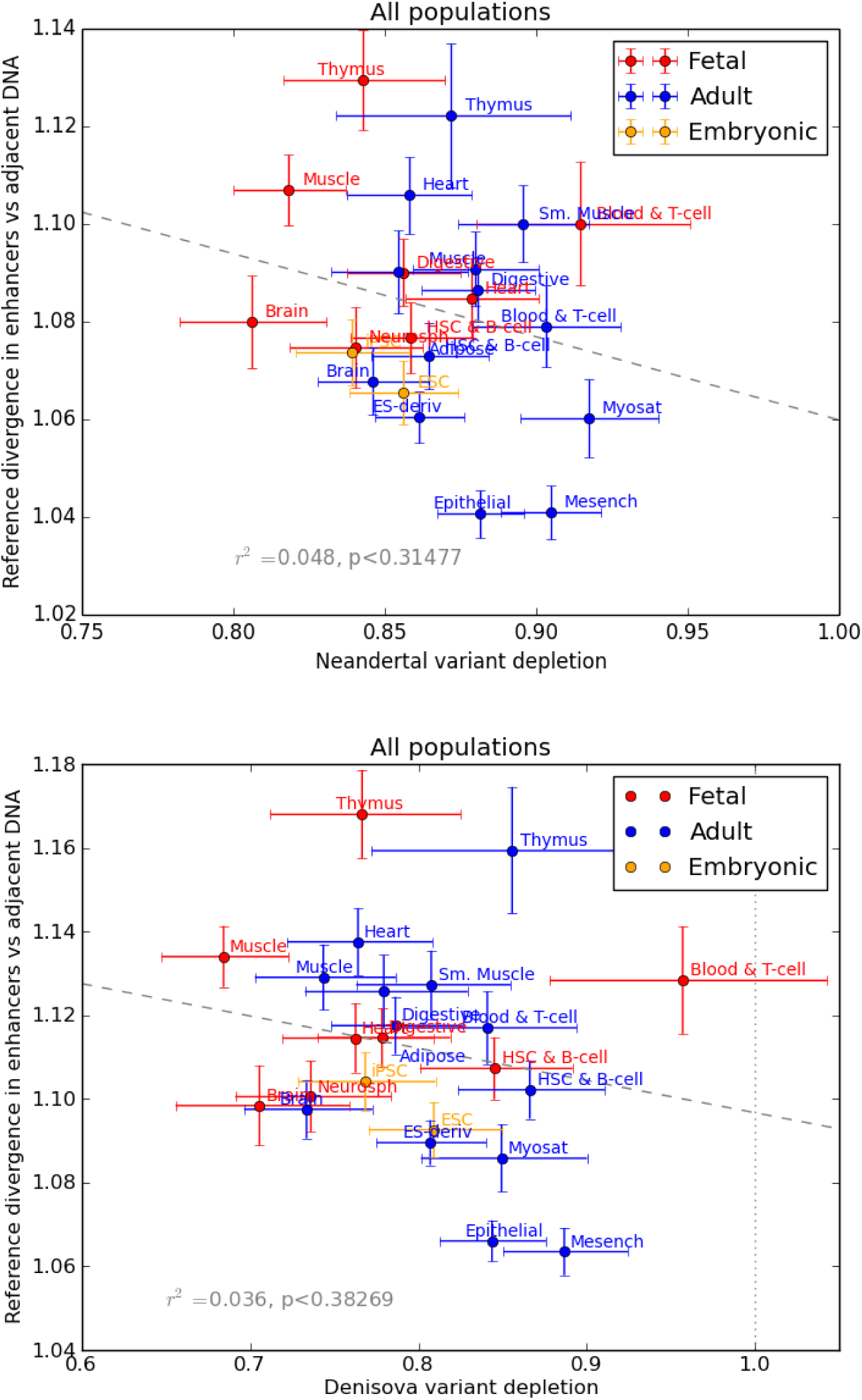
Within the set of enhancers active in each cell type, we measured divergence between human and archaic reference genomes and compared it to human/archaic divergence within adjacent matched control regions as described in Figure S2.1. The results are plotted here together with error bars derived from the binomial approximation to the Bernoulli distribution. We see no correlation across tissues between the amount of archaic divergence from the human reference and the degree of archaic allele depletion from enhancers. This absence of correlation suggests that differences between tissues in density of introgressed archaic variants are not driven by regional differences in the degree of divergence between archaic and human genomes.

**Figure S3.1.**
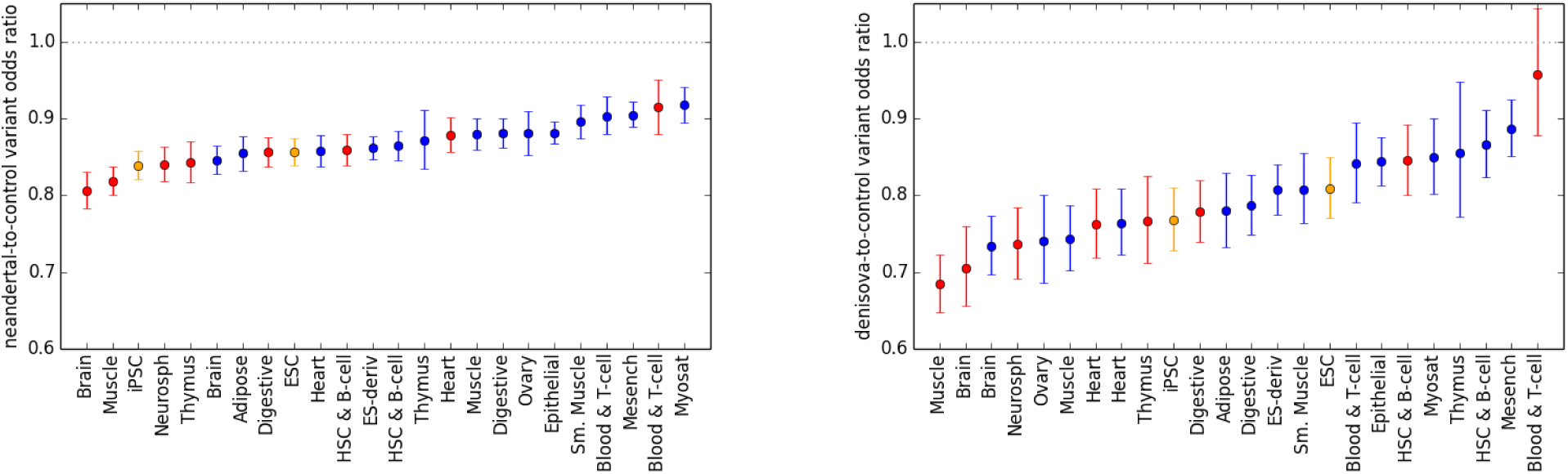
These plots show the data from Figure 3 with Neandertal and Denisovan odds ratios on separate plots for clarity.

**Figure S3.2.**
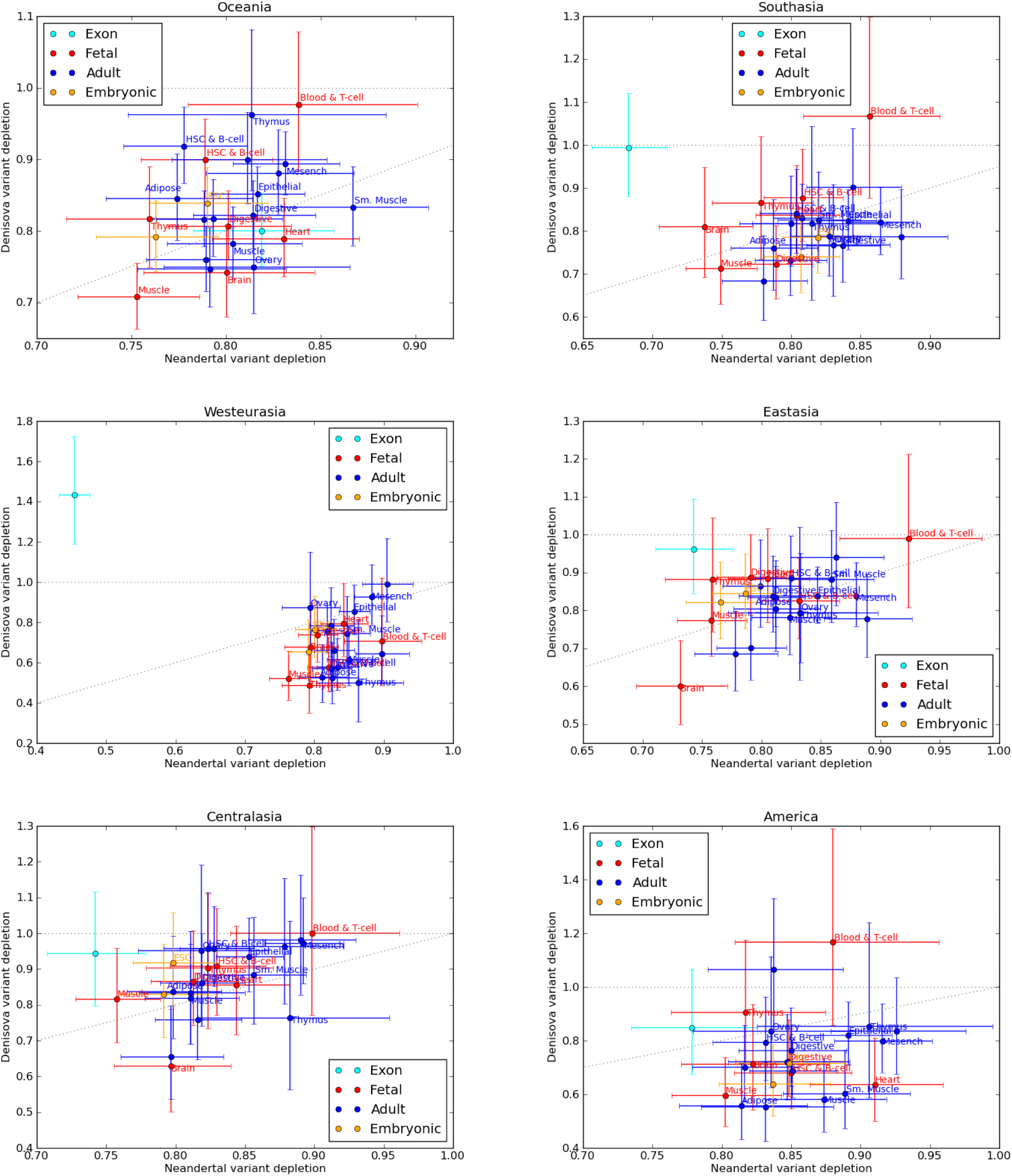
These plots show the joint distribution of Neandertal and Denisovan introgression depletion within each SGDP population separately. Although there are differences between populations, particularly since Denisovan introgression is sparse and noisy in general, all show that brain and fetal muscle enhancers are the most depleted of introgression. In most populations the blood & T-cell tissue is least depleted of introgression.

**Figure S4.1.**
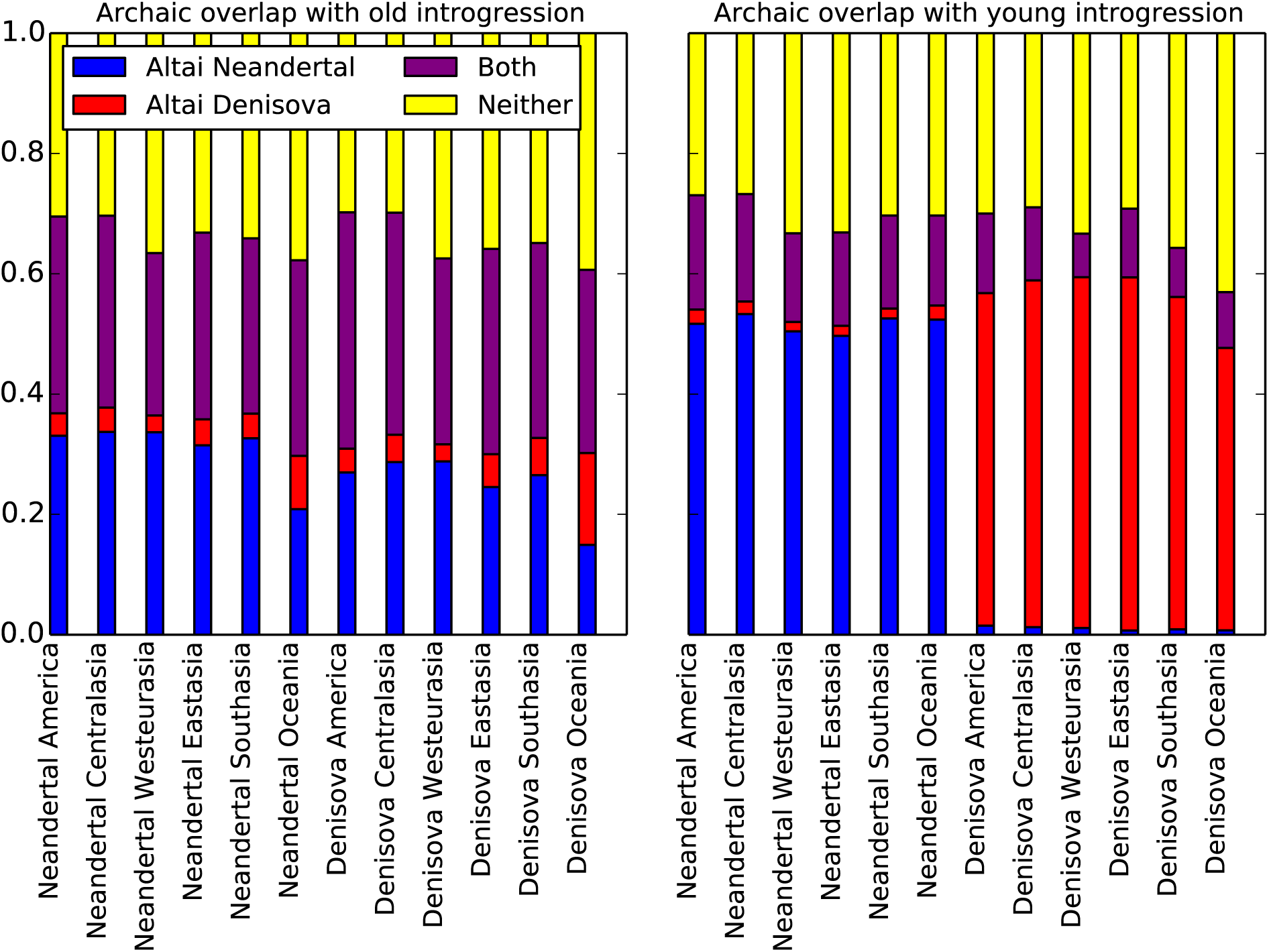
For each human SGDP population, this figure shows the fraction of introgressed nonreference alleles that are shared with the Altai Neandertal genome, the Altai Denisova genome, both, or neither. Recall that “young” introgression calls are SNPs that appear in call set 2 generated by Sankararaman, et al. while “old” calls appear in call set 1 for at least one archaic species but not in either set of young calls. In every population, 20% to 30% of old introgressed SNPs are shared with both archaic reference genomes, indicating that these alleles predate the divergence of Neandertals and Denisovans or are at least old enough to have passed between the two species by gene flow. In contrast, only 10% to 20% of young introgressed SNPs are present in both archaic genomes. In each set of young Neandertal introgression calls, over 45% of alleles are shared with the Altai Neandertal but not the Altai Denisovan; conversely, in each set of young Denisovan introgression calls, over 45% of alleles are shared with the Denisovans reference but not the Neandertal reference. Old Neandertal and Denisovan introgression calls look much more similar to each other: each contains 10% to 25% alleles found specifically in the Neandertal reference as well as 2% to 10% alleles found specifically in the Denisovan reference. These patterns of allele sharing support our hypothesis that the old calls are indeed older than the young calls.

**Figure S4.2.**
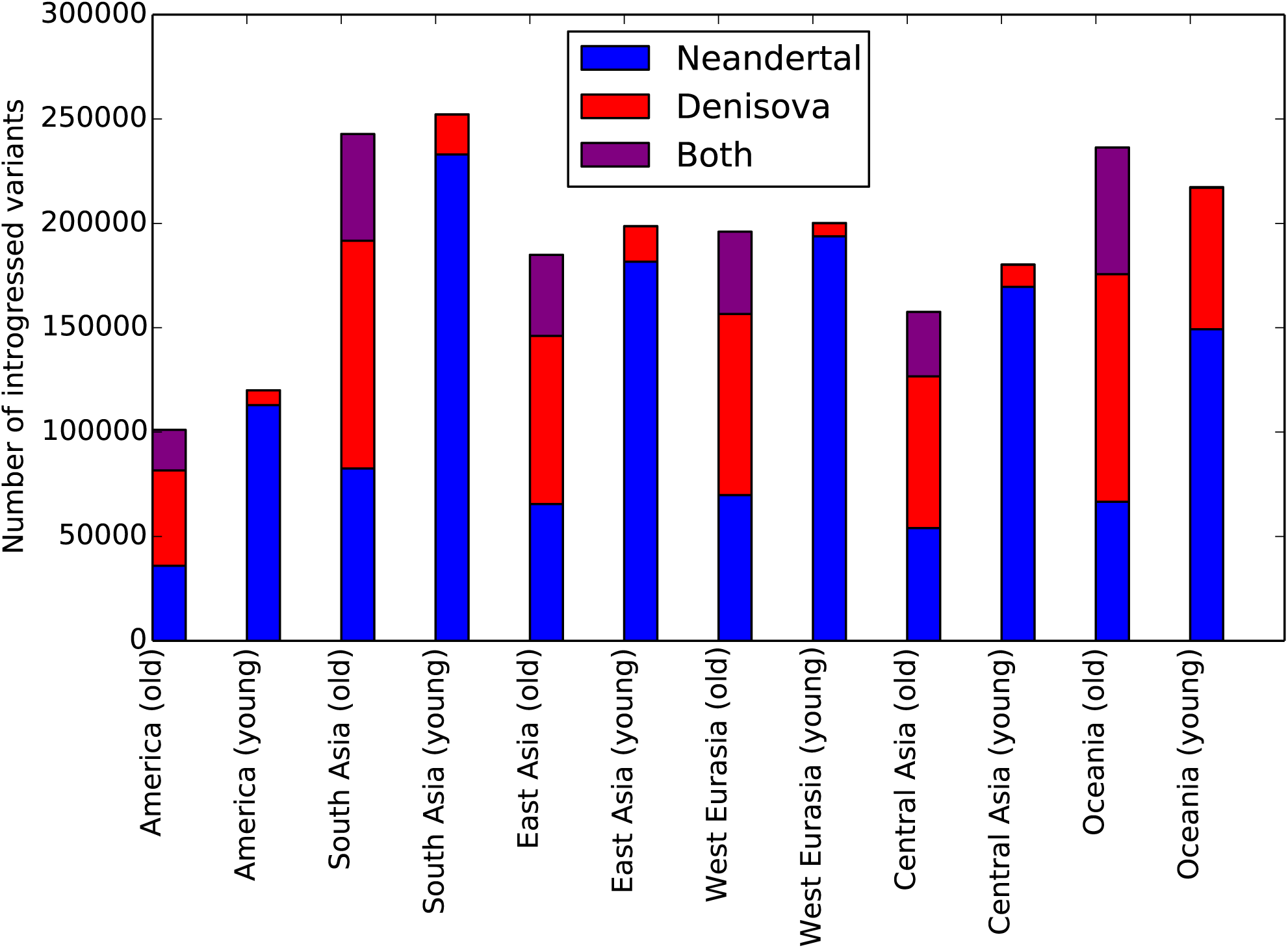
This chart shows the number of SNPs in each population that we classify as young versus old, based on their presence in the Sankararaman, et al. call sets 1 versus 2. Each SNP set is further subdivided into SNPs that appear in the Neandertal introgression call set only, the Denisova introgression call set only, or the intersection of both call sets.

**Figure S4.3.**
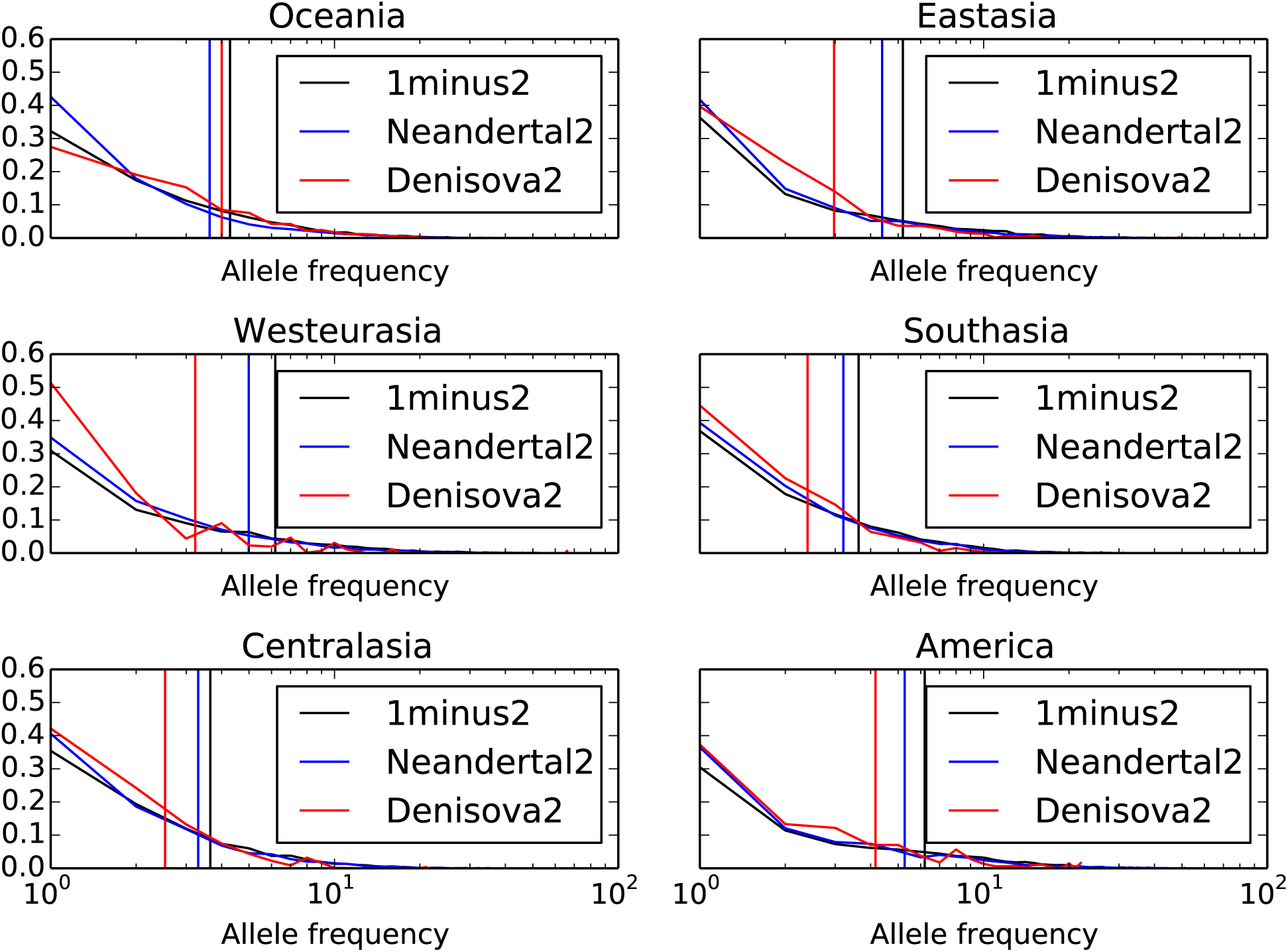
For each of the six SGDP populations, this figure shows the site frequency spectra of variants from the conditional random field call sets that we classify as young versus old. For each call set, the corresponding vertical line demarcates the mean allele frequency of that category. In each population, the old “1 minus 2” call set has the highest mean allele frequency, adding support to our hypothesis that these variants are older and/or less deleterious than either population-specific Call Set 2.

**Figure S5.1.**
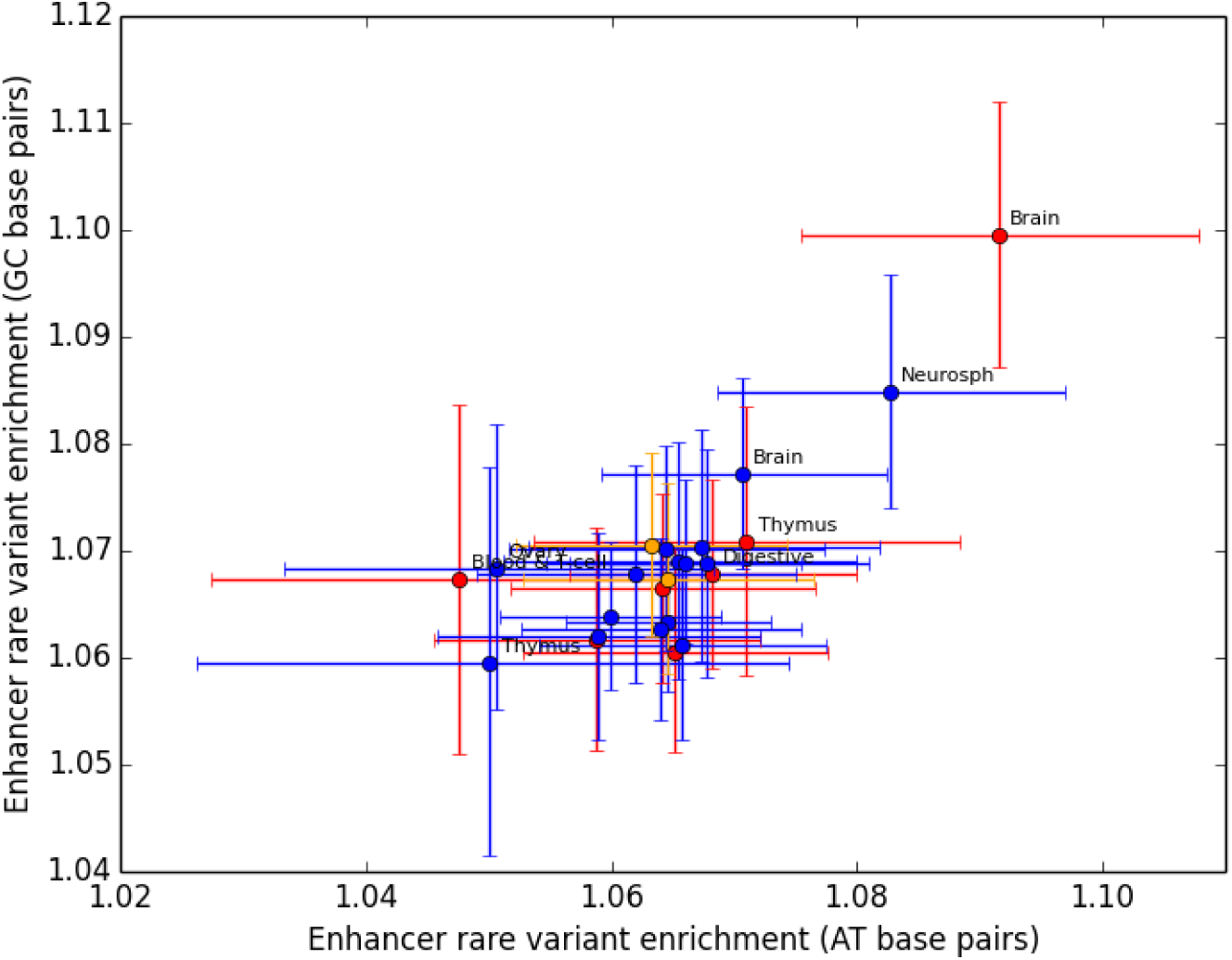
This figure was generated by partitioning the site frequency spectrum of each enhancer between SNPs that have GC ancestral alleles and SNPs that have AT ancestral alleles. The site frequency spectra of these two classes of sites are expected to be driven in opposite directions by GC biased gene conversion. However, the finding that brain enhancers are enriched for singletons holds up when we restrict to either GC-ancestral SNPs or AT-ancestral SNPs.

**Figure S5.2.**
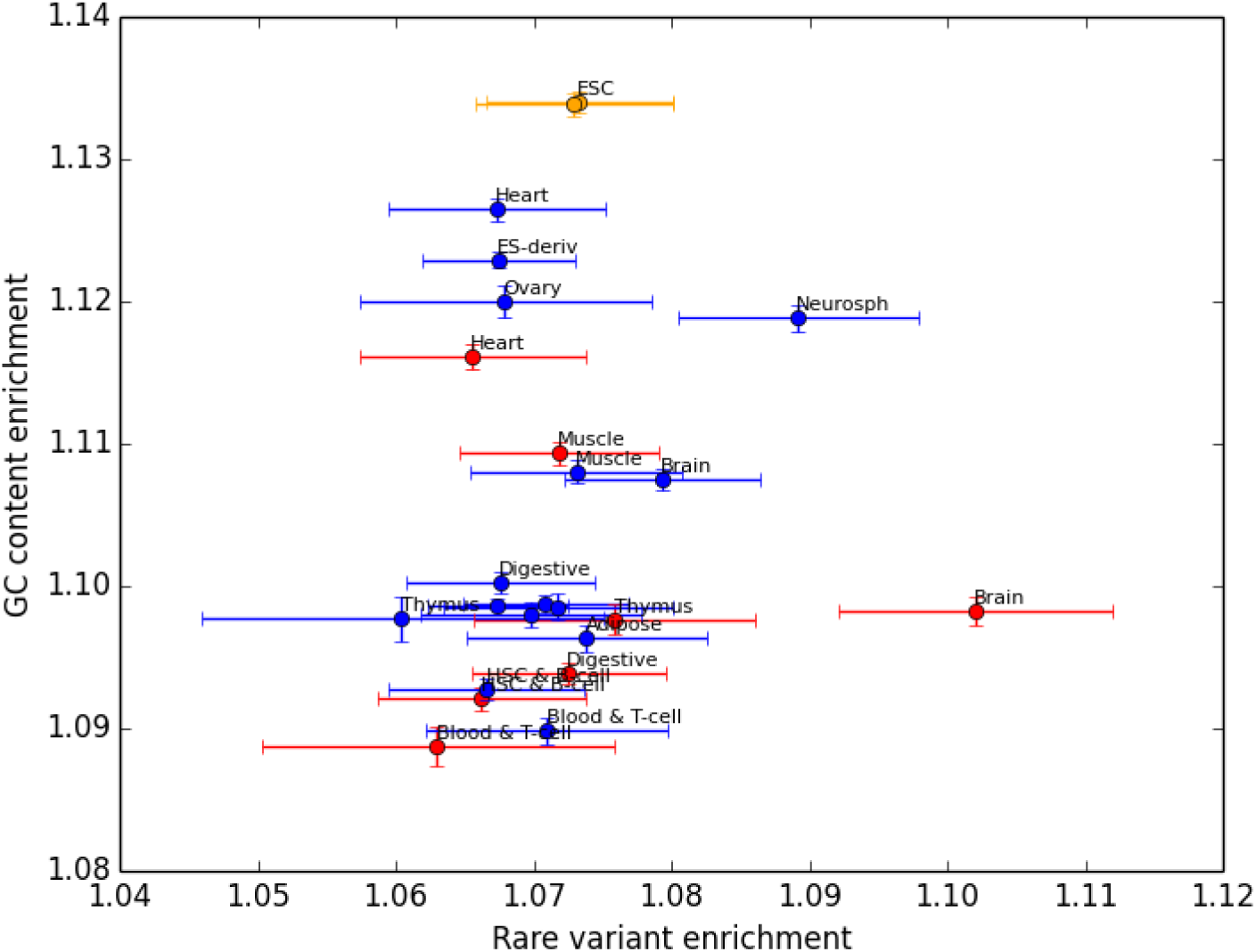
Enhancers are enriched for GC base pairs compared to adjacent genomic regions, and the degree of this enrichment varies between tissues. However, there is no correlation across tissues between GC content enrichment and the singleton enrichment that we attribute to purifying selection.

**Figure S5.3.**
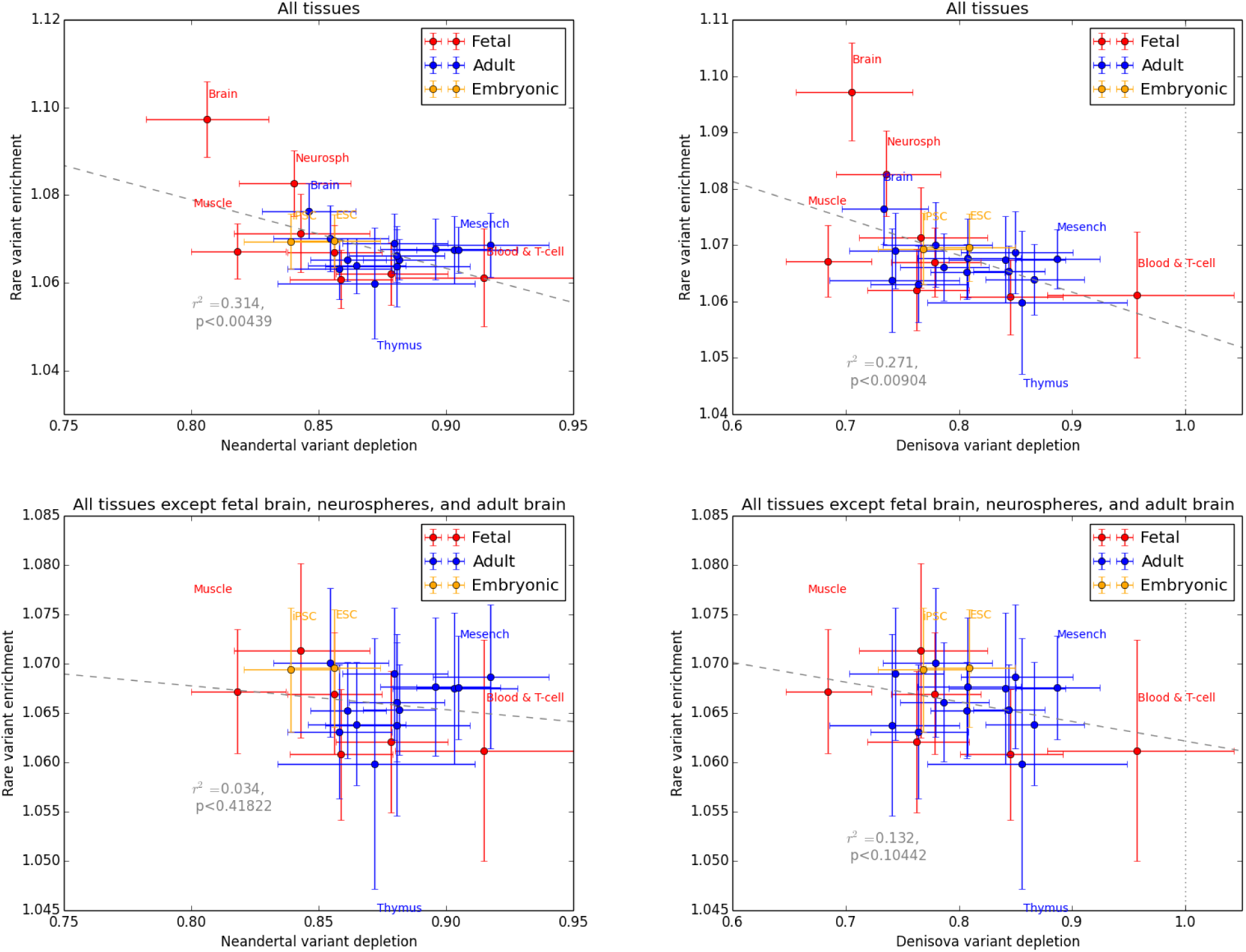
Although singleton enrichment is negatively correlated between tissues with both Neandertal and Denisovan variant depletion, the significance of this correlation disappears when all brain related tissues are excluded from the regression.

**Figure S5.4.**
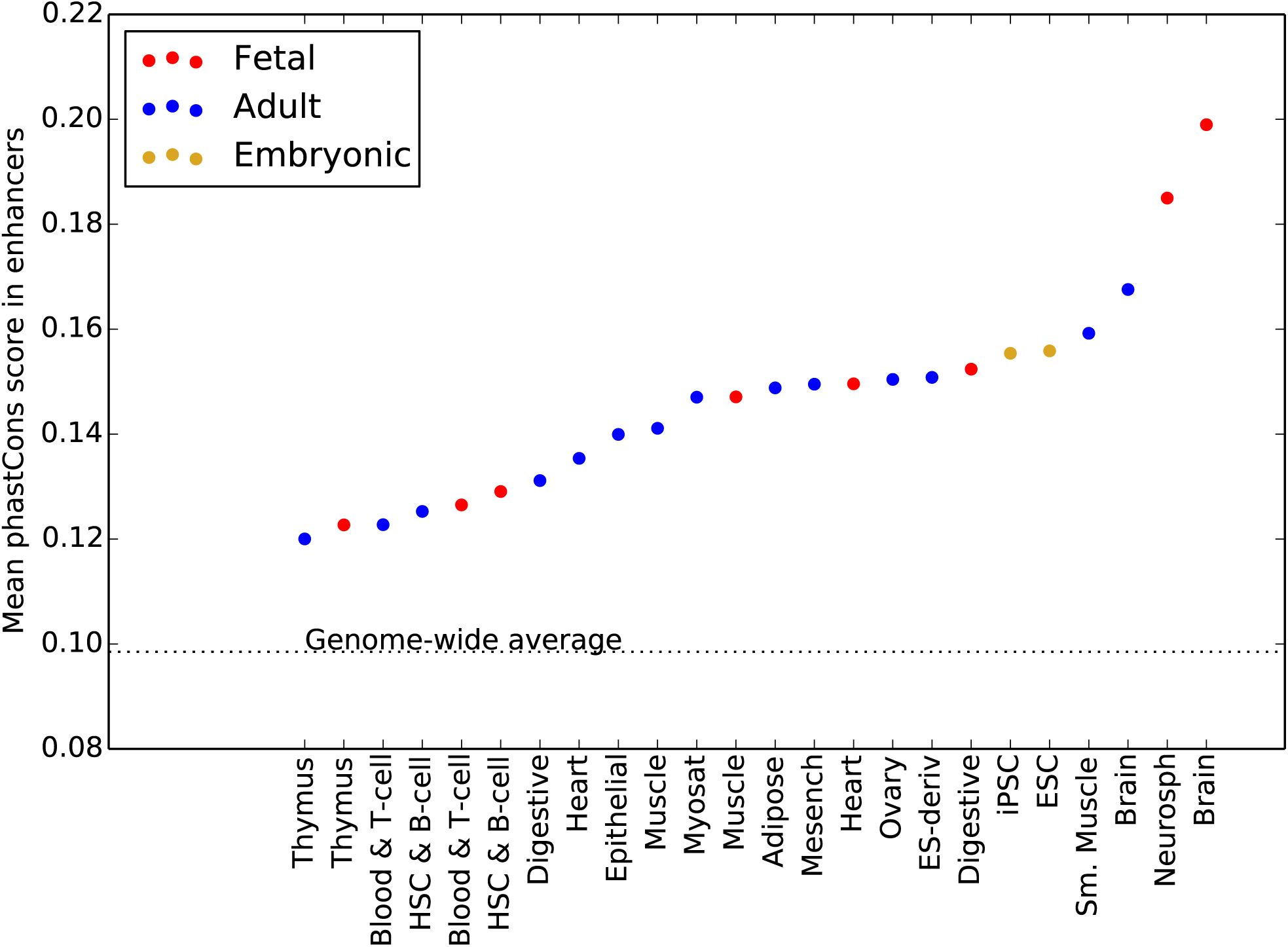
Across all tissues, enhancers have a mean phastCons score that is slightly elevated above the genomic mean, indicating that these regions are slightly conserved over phylogenetic timescales. Fetal brain and neurosphere enhancers have a higher mean phastCons score than enhancers active in any other tissues. This result mirrors our findings on the landscape of recent purifying selection as measured by site frequency skew: fetal brain enhancers are more conserved than other regulatory elements, but fetal muscle enhancers are not.

